# Lung and liver editing by lipid nanoparticle delivery of a stable CRISPR-Cas9 RNP

**DOI:** 10.1101/2023.11.15.566339

**Authors:** Kai Chen, Hesong Han, Sheng Zhao, Bryant Xu, Boyan Yin, Marena Trinidad, Benjamin W. Burgstone, Niren Murthy, Jennifer A. Doudna

## Abstract

Lipid nanoparticle (LNP) delivery of CRISPR ribonucleoproteins (RNPs) has the potential to enable high-efficiency *in vivo* genome editing with low toxicity and an easily manufactured technology, if RNP efficacy can be maintained during LNP production. In this study, we engineered a thermostable Cas9 from *Geobacillus stearothermophilus* (GeoCas9) using directed evolution to generate iGeoCas9 evolved variants capable of robust genome editing of cells and organs. iGeoCas9s were significantly better at editing cells than wild-type GeoCas9, with genome editing levels >100X greater than those induced by the native GeoCas9 enzyme. Furthermore, iGeoCas9 RNP:LNP complexes edited a variety of cell lines and induced homology-directed repair (HDR) in cells receiving co-delivered single-stranded DNA (ssDNA) templates. Using tissue-selective LNP formulations, we observed genome editing of 35‒56% efficiency in the liver or lungs of mice that received intravenous injections of iGeoCas9 RNP:LNPs. In particular, iGeoCas9 complexed to acid-degradable LNPs edited lung tissue *in vivo* with an average of 35% efficiency, a significant improvement over editing efficiencies observed previously using viral or non-viral delivery strategies. These results show that thermostable Cas9 RNP:LNP complexes are a powerful alternative to mRNA:LNP delivery vehicles, expanding the therapeutic potential of genome editing.

## Introduction

CRISPR-Cas9-based genome editing^1–3^ has the potential to provide wide-ranging genetic disease treatments^4–6^ if safe and effective methods for delivering CRISPR-based therapeutics can be developed^7,8^. Although viral delivery of CRISPR genome editors is the most widely used method for *in vivo* cell editing^9–11^, viral vectors can be immunogenic, carry the risk of vector genome integration and can induce off-target DNA damage due to continuous genome editor expression^12^. Alternative non-viral strategies for delivering CRISPR editors could address these limitations if issues of efficacy and toxicity can be overcome.

Lipid-nanoparticle (LNP):mRNA complexes are non-virally derived vehicles for *in vivo* delivery that have been remarkably successful at genome editing in the liver^13–15^. However, developing LNP:mRNA complexes that can edit non-liver tissues remains a challenge. Although LNPs can deliver mRNAs coding for Cre recombinase, luciferase and fluorescent proteins to non-liver organs, making the transition from reporter enzymes to CRISPR mRNA and sgRNA delivery has been inefficient^16,17^. LNP-mediated delivery of CRISPR mRNA and sgRNA faces challenges of sgRNA instability^18^, mRNA-mediated Toll-like receptor (TLR) activation^7^ and low translational efficiency of the large mRNAs encoding genome editors^19^. These challenges are inherent to the mRNA formulation but could be mitigated with alternative LNP delivery cargoes.

The direct delivery of genome editors in the form of RNP complexes^20^ has the potential to address several of the limitations associated with mRNA and viral-based delivery of CRISPR editors. In particular, RNPs are expected to elicit lower levels of TLR activation than mRNA and produce minimal off-target DNA damage due to their short intracellular half-life^21–23^. In addition, RNPs may offer higher *in vivo* editing efficiency compared to mRNA-based delivery methods by avoiding *in-situ* translation of large mRNA^19^ and providing natural protection of the sgRNA by high-affinity Cas9 binding^18^. Strategies for delivering RNPs include the use of complex polymers^24,25^, silica nanoparticles^26^, metal-organic frameworks^27^, LNPs^28–30^ and other formulations^31–33^. However, only LNPs have a proven track record of clinical use and established procedures for good manufacturing practice (GMP)^34^. A successful LNP-based delivery strategy for RNPs therefore has great translational potential. Nevertheless, RNPs lack the negative charge density needed for efficient LNP encapsulation. Furthermore, conditions to formulate LNPs usually consist of organic solvents that can denature proteins^13^. Although LNP-mediated delivery of SpyCas9 in the RNP format induced genome editing in the liver^30^, delivery to non-liver organs such as the lungs remains inefficient^17,29^.

We rationalized that the protein denaturation problem currently limiting LNP-based RNP delivery could be tackled using alternative, thermostable CRISPR enzymes. GeoCas9 has great potential for LNP-mediated delivery because of its higher thermal stability^35^ compared to commonly used editors such as *S. pyogenes* Cas9 (SpyCas9) or *Lachnospiraceae bacterium ND2006* Cas12a (LbCas12a). However, GeoCas9 has low genome editing efficiency and uses a large protospacer adjacent motif (PAM) that prevents it from editing a large fraction of the genome^35,36^.

In this study, we demonstrate that laboratory-evolved GeoCas9 mutants, iGeoCas9s, can edit mammalian cells with >100-fold higher efficiency than wild-type GeoCas9 and can edit cells and animal organs efficiently after LNP-mediated delivery. An LNP-based platform containing pH-sensitive PEGylated and cationic lipids enabled iGeoCas9-mediated editing of mouse neural progenitor cells (NPCs), human embryonic kidney 293T (HEK293T) cells and human bronchial epithelial (HBE) cells with efficiencies ranging from 4% to 99% depending on the locus. These iGeoCas9 RNP:LNPs could also induce homology-directed repair (HDR) upon co-delivery with ssDNA templates, as well as editing of mouse liver and lung tissues after intravenous injection. RNP:LNP formulations containing an acid-degradable PEGylated lipid edited an average of 56% of the entire liver issue in Ai9 mice, and formulations containing acid-degradable PEGylated and cationic lipids edited an average of 35% of the entire lung tissue. This level of lung editing observed with iGeoCas9 RNP:LNPs represents the most efficient lung editing observed to date using either viral or non-viral delivery strategies^37–39^. Collectively, these results demonstrate that evolved, thermostable genome editors delivered with LNPs can efficiently edit cells *in vitro* and *in vivo* and are a promising platform for developing CRISPR therapeutics.

## Results

### Directed evolution of GeoCas9 improves editing efficiency and PAM compatibility

GeoCas9 is a compact type II-C CRISPR-Cas9 protein that can function as a robust RNA-guided endonuclease at elevated temperatures (50‒65 °C is its optimal temperature range) or in the presence of human plasma^35^. These properties make GeoCas9 an attractive editor for delivery *in vivo*, particularly in the RNP format. However, GeoCas9 is far less effective than the canonical SpyCas9 at genome editing in mammalian cells and has a more restricted PAM. Wild-type GeoCas9 recognizes a PAM sequence of 5’-N_4_CRAA-3’ (where R is A/G) and can consequently target a much smaller fraction of the genome than SpyCas9, which has a PAM sequence of 5’-NGG-3’.

We rationalized that directed evolution could be used to improve the editing efficiency of GeoCas9 and also minimize its PAM sequence requirement. A bacterial dual-plasmid selection system^40–42^ was used to select for evolved active GeoCas9 variants, based on Cas9-mediated cleavage of a plasmid encoding the ccdB toxin gene under the control of an inducible pBAD promoter (**Fig. 1a**). To search for a reliable evolutionary starting point with minimal activity in the *E. coli* assay, we screened 20 different sgRNAs that target the ccdB gene at the protospacers associated with different PAM sequences (**Fig. S1**) and performed selection under two sets of conditions (37 °C or 30 °C for 1.5 hours). Target sequence #6 with a disfavored PAM sequence (ggat<u>GAAA</u>) gave a minimal survival rate under either condition (<0.1% for 30 °C and 2-5% for 37 °C) and was chosen for engineering. Libraries of GeoCas9 mutants were generated by targeting different domains of the protein for random mutagenesis (**Fig. S1**) and then subjected to the selection system under the conditions at 30 °C. To amplify the most active mutants in these libraries, selected mutants were collected and subjected to another round of selection (**Fig. S1**). Sequencing of the selected colonies identified frequently appearing beneficial mutations from each library (**Fig. S1**). For instance, the library targeting BH + Rec domains for random mutagenesis generated mutant GeoCas9(R1) bearing four mutations, E149G, T182I, N206D and P466Q, which gave >95% survival (vs. <5% with the wild-type protein) in the bacterial assay. Addition of further beneficial mutations identified in the library targeting RuvC + HNH + WED domains, including E843K, K908R, E884G and Q817R, to the mutant R1 construct produced a new lineage of variant GeoCas9 proteins (**Fig. 1b**). Combining a total of eight beneficial mutations yielded a composite mutant, GeoCas9(R1W1), which possesses greatly improved target dsDNA cleavage activity (**Fig. S1**) and well-preserved thermostability (T_m_: 55 °C *vs.* 60 °C, R1W1 mutant *vs.* wild-type protein) (**Fig. 1c**; **Fig. S2**).

**Figure 1.**
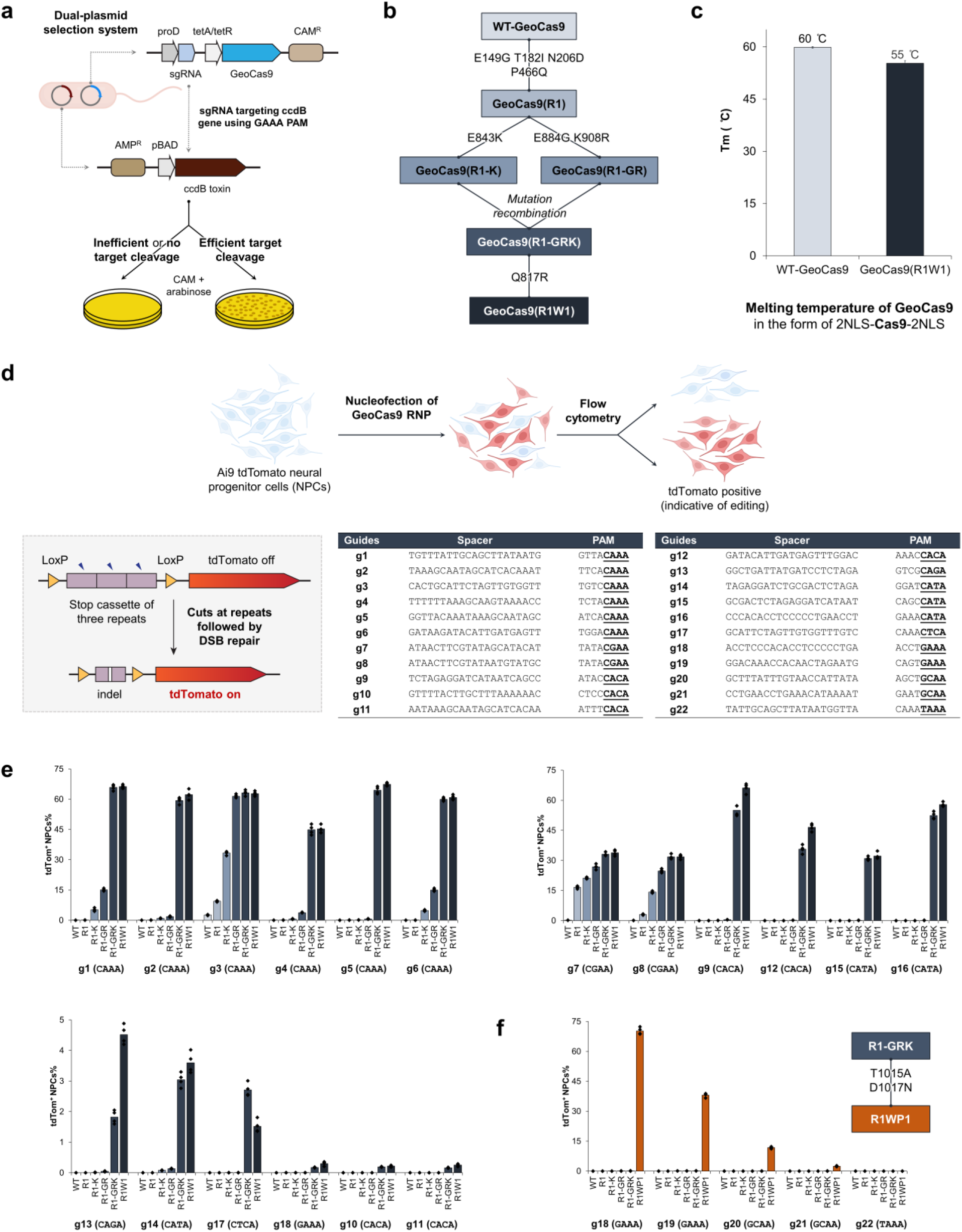
Directed evolution of GeoCas9 improves its editing efficiency by orders of magnitude and broadens its PAM compatibility. **a.** Schematic diagram of the direct evolution system used to evolve GeoCas9, based on bacterial selection. **b.** Evolutionary lineage of GeoCas9 mutants. **c.** Evolved GeoCas9 (mutant R1W1) has a higher melting temperature than wild-type GeoCas9. Melting temperature measurements of WT and engineered GeoCas9 proteins were measured via a thermal shift assay. **d.** Schematic diagram of GeoCas9-mediated genome editing of NPCs isolated from Ai9 mice. The spacer and PAM sequences of the GeoCas9 gRNAs were designed to turn tdTomato if successful editing occurs. **e.** GeoCas9 mutants edit NPCs with significantly higher efficiency than wild-type GeoCas9 after electroporation-mediated delivery. Genome editing efficiencies quantified based on tdTom^+^ signals with the whole lineage of GeoCas9 mutants paired with different sgRNAs. **f.** Engineering of GeoCa9 can broaden its PAM specificity. n = 4 for each group, mean ± s.e.m.

The genome editing ability of the engineered GeoCas9 mutants was assessed in neural progenitor cells (NPCs) isolated from Ai9 tdTomato mice. In these cells, successful editing of a stop cassette sequence turns on tdTomato gene expression (**Fig. 1d**). Twenty-two sgRNAs were designed to target the SV40-derived poly(A) region using various PAM sequences. RNPs assembled from GeoCas9 mutants and these individual sgRNAs were electroporated into NPCs, and the percentage of tdTomato-positive cells was determined by flow cytometry. The evolved mutant, GeoCas9(R1W1), edited cells with >100-fold greater efficiency relative to the wild-type GeoCas9 with most sgRNAs investigated (**Fig. 1e**). In addition to editing NPCs, the evolved R1W1 mutant also exhibited robust genome editing in human embryonic kidney (HEK293T) cells and was able to reduce expression of enhanced green fluorescent protein (EGFP) with up to 99% editing efficiency (**Fig. S3**). These experiments demonstrate that the engineered GeoCas9(R1W1) can accept a broader range of PAM sequences, including but not limited to 5’-N_4_CNNA-3’ (vs. wild-type PAM sequences: 5’-N_4_CRAA-3’) (hereafter GeoCas9(R1W1) is referred to as iGeoCas9(C) for improved GeoCas9 targeting <u>C</u>-based PAM sequences).

To further expand the PAM compatibility of the engineered GeoCas9, the mutations T1015A and D1017N identified from the library targeting the WED + PI domains (**Fig. S1**) were incorporated into a later variant in the engineering lineage, GeoCas9(R1-GRK), to create GeoCas9(R1WP1) (hereafter referred to as iGeoCas9(G)) that alters the preference of the first base in the essential 4-nt PAM sequence from C to G (**Fig. 1f**). Taken together, these results show that directed evolution can be used to engineer GeoCas9 for improved genome editing activity and broadened PAM compatibility.

### iGeoCas9 RNP formulated in LNPs efficiently edits cells in vitro

The engineered iGeoCas9s have the potential to induce genome editing in cells and tissues that are not readily editable by other enzymes due to poor stability and/or limited delivery efficiency^17,29^. To test this, we compared the editing activity of iGeoCas9(C) to that of two established genome editors, SpyCas9 and iCas12a, an engineered version of LbCas12a^42^ (**Figs. 2a, S4**). Delivery by RNP nucleofection showed that all three of these enzymes generated robust and similar levels of genome editing in tdTomato NPCs. However, delivery of these RNPs using LNPs produced markedly different results: iGeoCas9 RNP:LNP delivery resulted in >2-fold higher editing efficiency compared to SpyCas9 RNP:LNPs, and iCas12a RNP:LNP delivery did not produce detectable editing in these cells. The improved performance of iGeoCas9 RNP relative to other CRISPR-Cas9 RNPs could be due to its higher stability and thus higher specific activity per LNP (**Fig. S4**, **S5**)^13^. In addition, the larger size of the sgRNA for iGeoCas9 compared to SpyCas9 (139 versus 96 nucleotides) generates an RNP with increased negative charges, which could facilitate LNP encapsulation (**Fig. 2a**).

**Figure 2.**
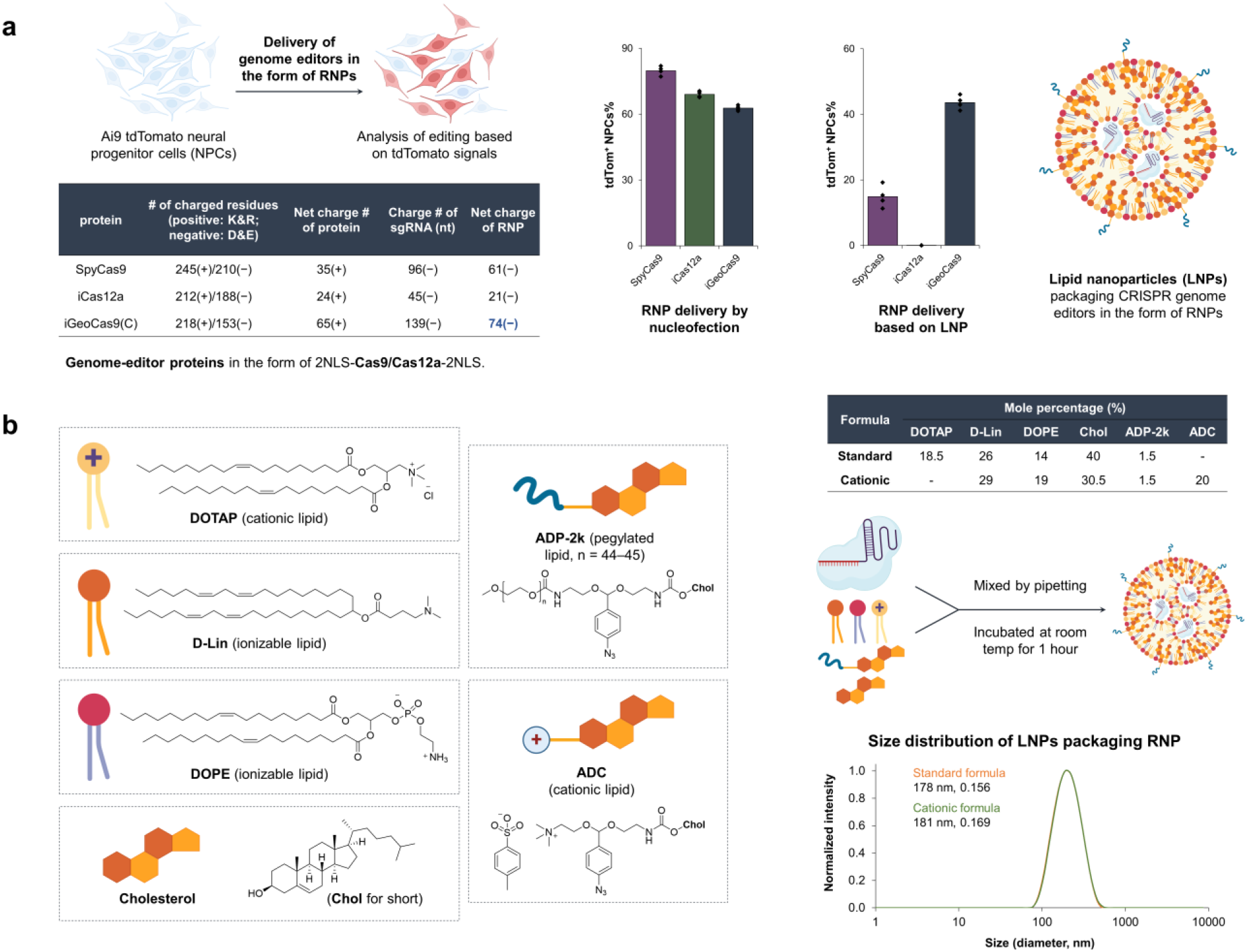
iGeoCas9 edits cells more efficiently than SpyCas9 or iCas12a after LNP-mediated delivery. **a.** iGeoCas9, SpyCas9, and iCas12a edit cells with similar efficiency after nucleofection. However, iGeoCas9 edits cells more efficiently after LNP-mediated delivery than either SpyCas9 or iCas12a, n = 4 for each group, mean ± s.e.m. **b.** Chemical structures of the lipids used in this study, two formulations were identified that delivered iGeoCas9 RNP efficiently, termed standard and cationic (see table for details). Dynamic light scattering (DLS) of standard and cationic LNPs demonstrate they have sizes of 178 nm and 181 nm.

To set up a robust LNP-based system for iGeoCas9 RNP delivery, we further optimized the lipid formulation for RNP encapsulation and LNP assembly. We used four commercial lipids, including 1,2-dioleoyl-3-trimethylammonium propane (DOTAP), (6Z,9Z,28Z,31Z)-heptatriaconta-6,9,28,31-tetraen-19-yl 4-(dimethylamino)butanoate (D-Lin), dioleoyl-sn-glycero-3-phosphoethanolamine (DOPE), cholesterol, and two synthetic lipids derived from cholesterol, ADP-2k and ADC, which are newly developed for mRNA delivery^43^ (**Fig. 2b**). The pegylated lipid, ADP-2k, proved to be key to the successful encapsulation of RNPs into LNPs and delivery to NPCs (**Fig. S4**). Low percentages (<1%) of ADP-2k led to relatively large particle sizes, which was not beneficial for LNP delivery; on the other hand, high percentages (>5%) of ADP-2k resulted in smaller particle size but also reduced the efficacy of RNP packaging as measured by reduced editing in NPCs. These observations correspond to the known behaviors of pegylated lipids in enhancing LNP stability, controlling particle size and regulating circulation time^44,45^.

We next examined several pegylated lipids, commercial and synthetic, for their ability to encapsulate and deliver iGeoCas9 RNPs in LNPs (**Fig. S4**). The commonly used 1,2-dimyristoyl-rac-glycerol-methoxy(poly(ethylene glycol)) (DMG-PEG) and other PEG lipids derived from DOPE were found to be detrimental to NPC viability, presumably due to the surfactant properties of PEG-lipids^46^. Interestingly, the synthetic pegylated lipid ADP-2k exhibited minimal toxicity in NPCs, and with its inclusion in LNPs, we observed >90% cell viability. Two dipeptide-fused PEG lipids, Pep-1k and Pep-2k, also showed good biocompatibility and high editing levels in NPCs (**Figs. S4**). The reduced toxicity and enhanced delivery efficiency of these three PEGylated lipids, ADP-2k, Pep-1k and Pep-2k, stem from the pH-sensitive, acid-degradable acetal linker used in their synthesis (**Fig. S6**). Specifically, the labile acetal linker is cleaved in the late endosome stage of LNP delivery at a pH of 5−6, which frees the PEG moiety from the lipid molecule to reduce cytotoxicity while destabilizing the endosome to promote RNP release into the cytosol (**Fig. S6**)^43^. Further optimization of other parameters of LNP assembly (including mole and volume ratios of lipids to RNP, and salt concentration in the buffer, **Fig. S7**) established two sets of lipid formulations, a standard formula (with DOTAP as the cationic lipid) and a cationic formula (with ADC as the cationic lipid). Both of these formulations can encapsulate iGeoCas9 RNPs to produce good particle uniformity (diameter: 170−180 nm, PDI: 0.13−0.17), an important feature of efficacious LNPs^47^ (**Fig. 2b**).

The genome editing efficacy of iGeoCas9 RNP:LNP complexes were evaluated in NPCs by targeting the SV40-derived poly(A) stop cassette to turn on tdTomato. RNPs were assembled using the engineered iGeoCas9 mutants with corresponding sgRNAs and then encapsulated into LNPs of the standard formulation. Quantification of genome editing by sorting tdTomato-expressing cells after LNP treatment established that LNP-based delivery had comparable delivery efficacy to nucleofection (**Fig. 3a**). We next tested whether changes to the sgRNA could further enhance editing efficiency using the LNP delivery strategy. We extended the protospacer region from 21nt to 23 or 24nt and introduced 2’-*O* methylation and phosphorothioate linkages to the last three nucleotides at both the 5’- and 3’-ends. These chemical modifications, which enhance the chemical stability of the sgRNA^18^, are beneficial to RNP delivery with LNPs (**Fig. 3b**). The LNP strategy was also capable of delivering iGeoCas9 RNPs to HEK293T cells and disrupting expression of an EGFP transgene with comparable efficiency to that observed using nucleofection (**Fig. 3c**). The cationic lipid formulation for LNP assembly was found to be slightly more effective for RNP delivery to HEK cells. Together these experiments establish a robust LNP-based system for delivering iGeoCas9 RNPs to different cell lines to perform effective genome editing.

**Figure 3.**
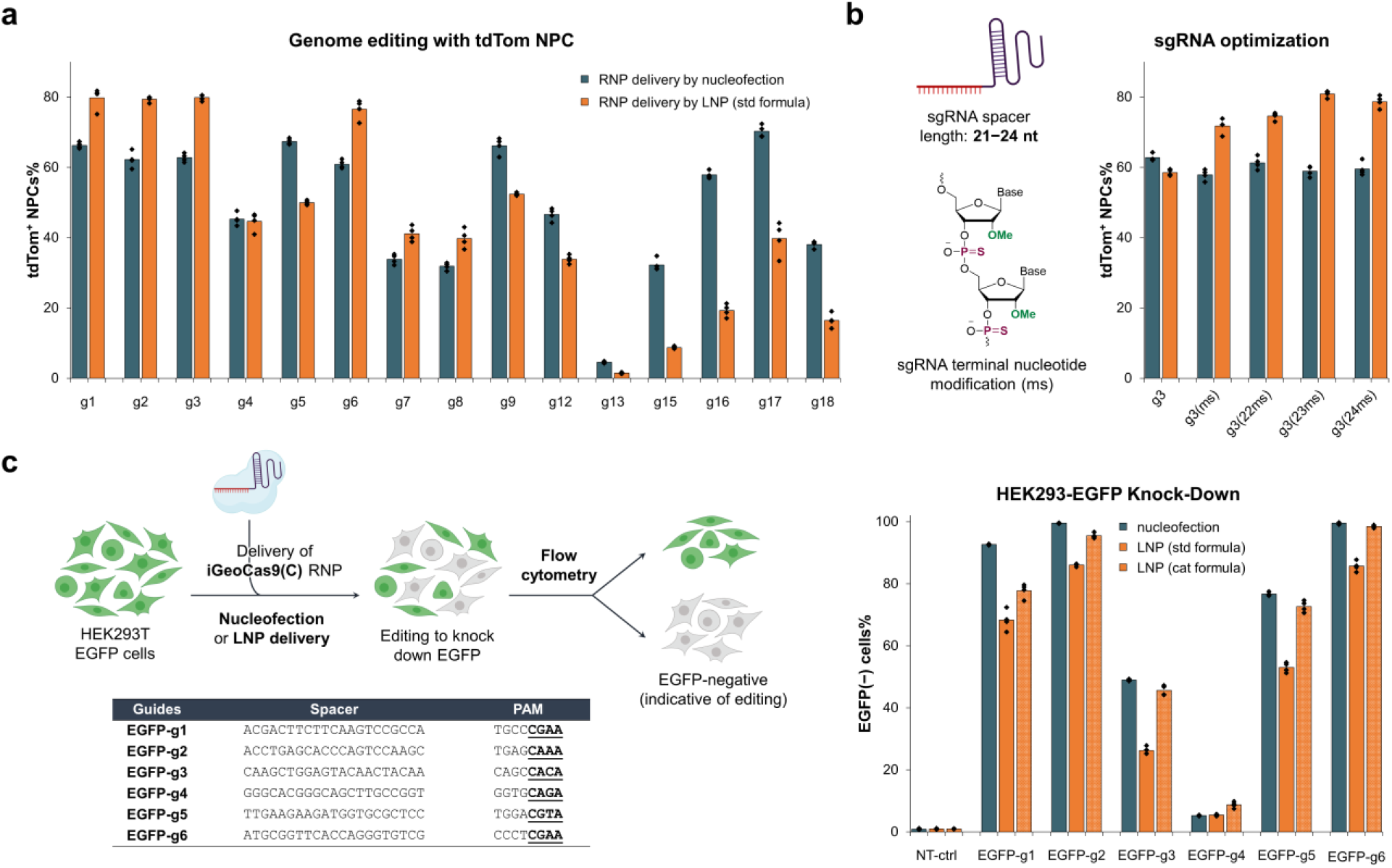
iGeoCas9 RNP:LNP complexes can edit a wide range of genomic targets and multiple different cell lines. **a.** LNP-mediated delivery of iGeoCas9 RNPs edits NPCs with an efficiency comparable to nucleofection. **b.** Chemical modification of the iGeoCas9 sgRNA improves its editing efficiency after LNP-mediated delivery (ms = methoxy and phosphorothioate). **c.** Comparison of the genome editing levels in HEK293T cells based on nucleofection and LNP-assisted delivery of iGeoCas9 RNP. (Left) Schematic diagram of iGeoCas9-mediated genome editing of EGFP HEK293T cells, resulting in the knock-out of EGFP fluorescence. (Right) Genome editing efficiencies quantified based on EGFP(‒) signals using the engineered GeoCas9 paired with different sgRNAs. n = 4 for each group, mean ± s.e.m.

### iGeoCas9 RNPs delivered with ssDNA templates induce site-specific integrations in cells

We next tested whether LNPs can co-deliver iGeoCas9 RNPs with a ssDNA template to induce site-specific genomic integrations through homology-directed repair (HDR). We first characterized the physical features of LNPs co-packaging iGeoCas9 RNPs and ssDNA templates of 180−200nt in length (**Fig. 4a**). Interestingly, in the presence of ssDNA (with a mole ratio of 1:1 for RNP:ssDNA), the nanoparticle size was reduced from ∼180 nm to 140−150 nm. This phenomenon is consistent with a recent study showing that ssDNA helps RNP encapsulation into LNPs and prevents LNP aggregation by transient binding to Cas9 RNPs^30^.

**Figure 4.**
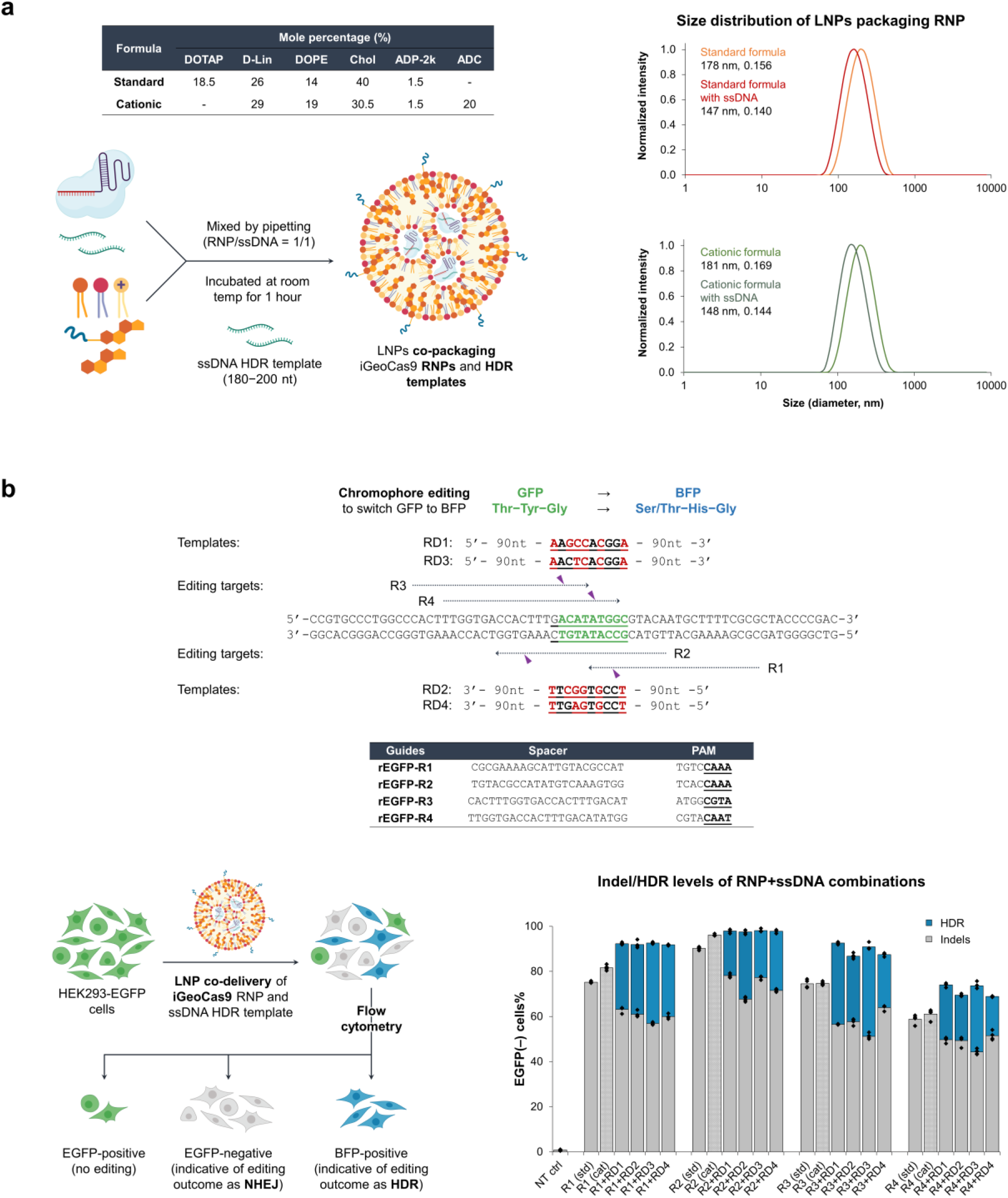
Co-delivery of iGeoCas9 RNPs and ssDNA templates with LNPs efficiently generates HDR in cells. **a.** Characterization of LNPs encapsulating iGeoCas9 RNPs and ssDNA templates. **b.** Co-delivery of iGeoCas9 RNPs and ssDNA HDR templates with LNPs edits the chromophore of EGFP to BFP in HEK293T cells. (Upper) Target and donor designs for GeoCas9-mediated chromophore editing. (Lower) Genome editing efficiencies quantified based on EGFP/BFP signals using the engineered iGeoCas9 (in the form of NLS-iGeoCas9(C)-2NLS) paired with different sgRNAs ± ssDNA templates. iGeoCas9 RNP = ss DNA generates between 20-40% HDR in HEK293T cells. n = 4 for each group, mean ± s.e.m.

We investigated whether the co-delivery of iGeoCas9 RNPs and ssDNA templates in LNPs could switch EGFP to the blue fluorescent protein (BFP) in a model HEK293T cell line (**Fig. 4b**). In this cell-based assay, editing of the chromophore Thr-Tyr-Gly in the EGFP transgene by HDR installs a Ser/Thr-His-Gly chromophore and converts EGFP into BFP. Four sgRNAs, rEGFP-R1 to -R4, were designed to target the coding and non-coding strands in the chromophore region for editing. To avoid possible recutting events after incorporation of the desired edits, four ssDNA HDR templates were designed to introduce GFP-to-BFP edits together with additional silent mutations in the DNA sequence. BFP signals were observed with the co-delivery tests based on all 16 combinations of RNPs and ssDNA templates using the standard lipid formulation for LNP assembly. HDR levels indicated by the percentage of BFP-positive cells were quantified by flow cytometry, ranging from 20% to 40% based on different RNP + ssDNA combinations, while 50−75% editing outcomes were determined as non-homologous end-joining (NHEJ), indicated by **Figs. 4b**, **S8**). Interestingly, the HDR experiments produced overall higher editing (HDR + NHEJ) levels compared to EGFP knockdown by RNP only, consistent with the role of ssDNA in promoting RNP encapsulation into LNPs. We wondered whether other anionic polymers, such as poly-L-glutamate (MW 15−50kDa) and heparin (MW 10−30kDa), could have similar effects (**Fig. S9**). As expected, the anionic polymer poly-L-glutamate also reduced the LNP size and modestly improved editing levels. However, the addition of heparin resulted in reduced editing, probably due to its inhibitory effect on Cas9 function. These results suggest that anionic polymers promote RNP packaging into LNPs through the charge interaction between the polymer additives and cationic lipids (**Figs. S5**, **S9**).

LNP-based co-delivery of iGeoCas9 RNPs and ssDNA templates was further used to induce HDR at endogenous genomic sites in human cells. Four sets of guide RNAs and corresponding donor ssDNAs were designed to target different sites in the EMX1 gene and AAVS1 locus, respectively, for genome editing based on HDR (**Fig. 5a**). Both the standard and cationic LNP formulations were evaluated for their ability to deliver editing materials to HEK293T cells. HDR levels were quantified using next-generation sequencing (NGS), and LNP:RNP:ssDNA complexes generated up to 66% HDR, with total editing levels up to 95%. We then applied this co-delivery system to cell lines of disease models and tested whether the LNP-based editing materials can correct pathogenic mutations. Cystic fibrosis is a genetic disease caused by mutations in the CFTR gene, which encodes the ion channel protein cystic fibrosis transmembrane conductance regulator (CFTR). Two human bronchial epithelial cell lines (16HBEge) containing nonsense mutations in the CFTR gene, G542X and W1282X, respectively, were employed for the HDR tests (**Fig. 5b**). iGeoCas9 RNPs and HDR donors were co-delivered to the HBE cells, resulting in 7% HDR that reverted the pathogenic mutations G542X and W1282X, as quantified by NGS. These results suggest that further development of LNP-based RNP delivery may have therapeutic utility for restorative genome editing in the future.

**Figure 5.**
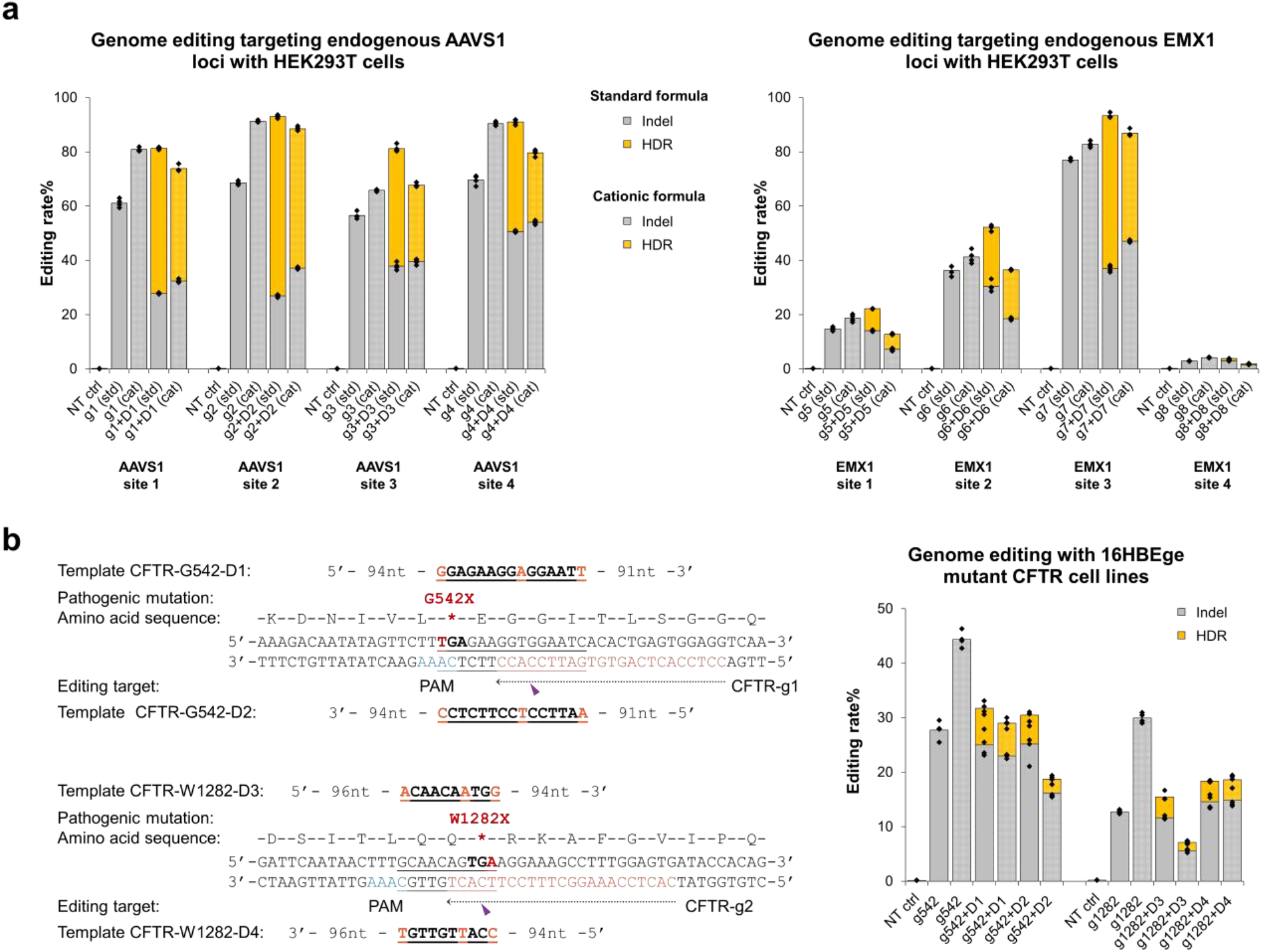
LNP-based delivery of iGeoCas9 RNP and ssDNA templates edits endogenous sites of the human genome. **a.** Genome editing efficiencies (indels and HDR) by iGeoCas9 paired with different sgRNAs ± ssDNA templates, as quantified by NGS. **b.** Editing of pathogenic mutations in the CFTR gene through HDR. (Left) Target and donor designs for iGeoCas9(C)-mediated editing of pathogenic mutations. (Right) Genome editing efficiencies quantified by NGS. n = 4 for each group, mean ± s.e.m. iGeoCas9 used here is in the form of NLS-iGeoCas9(C)-2NLS.

### LNP-based delivery of iGeoCas9 RNPs edits multiple organs efficiently in vivo

Having demonstrated that iGeoCas9 RNPs can be delivered by LNPs to a variety of cell lines for different editing purposes, we then asked if the RNP:LNP strategy can induce *in vivo* genome editing in mice following intravenous injection. More importantly, we wanted to test whether iGeoCas9 RNPs can be delivered to organs beyond the liver, which represents a major challenge for LNP-mediated delivery of CRISPR genome editors and other molecular cargoes.

We employed tdTomato Ai9 mice to assess the delivery and editing efficacy *in vivo* using our LNP-based delivery system for iGeoCas9 RNPs (**Fig. 6a**). The success of organ-specific mRNA delivery using SORT LNPs^13^ prompted us to test the ability of different lipid formulations to deliver genome editors to organs beyond the liver. Due to the observed benefit of ssDNA on RNP encapsulation and LNP delivery^30^, a 200-nt ssDNA was used to enhance iGeoCas9 RNP encapsulation in LNPs prepared with different lipid formulations. We performed two retro-orbital injections of LNPs at a dose of 3 mg/kg based on sgRNA (∼1 nmol RNP/injection based on mouse weight of 18−20 grams) at 3-day intervals. Mice were sacrificed a week after the second injection, and organs, including the liver, lung, spleen, heart and kidney, were collected to analyze the tdTomato signal as a reporter of genome editing outcome (**Fig. 6b**). Imaging of the organ slices together with flow quantification of tdTomato-positive cells revealed that iGeoCas9 RNPs can be delivered and induce robust genome editing *in vivo* with up to 56% editing in the liver and 35% editing in the lungs, as quantified by flow cytometry of the dissociated corresponding tissue (n = 6, controls were PBS-treated Ai9 mice) (**Figs. 6c, S10, S11**). Notably, our standard LNP formulation drove the delivery of RNPs primarily to the liver, while a modified cationic formulation using 40% ADC as the cationic lipid shifted the delivery specificity to the lungs with unprecedented genome editing levels. In addition, genome editing was also observed in other tissues known as challenging delivery targets, such as the heart (0.2% genome editing indicated by tdTomato expression level, **Fig. S11**). Future improvements to editing in the heart and other organs could expand the therapeutic utility of LNP-based delivery of genome editing RNPs.

**Figure 6.**
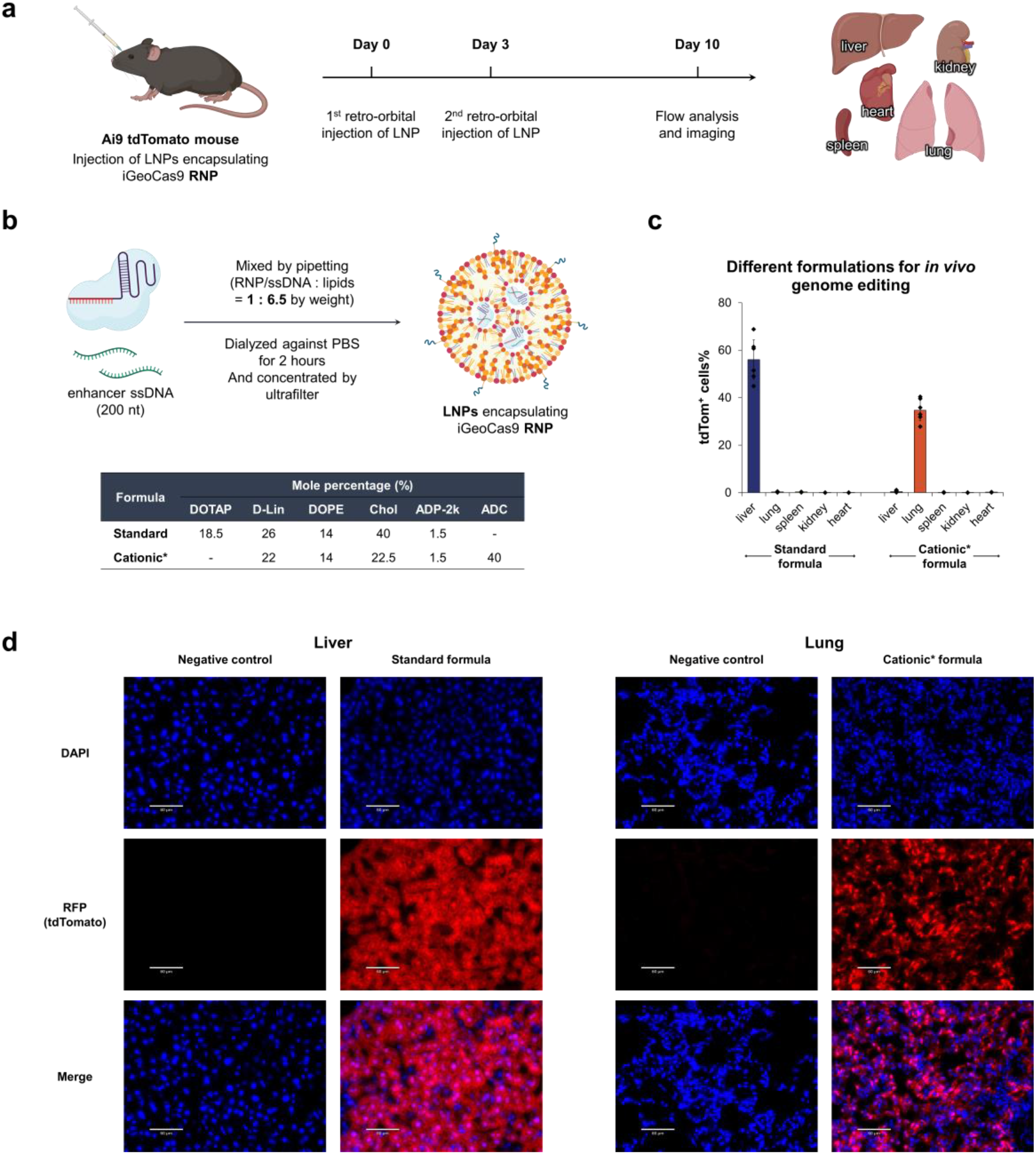
iGeoCas9 RNP:LNPs efficiently edit the liver and lungs of mice. **a.** Schematic diagram of the experimental design used to evaluate iGeoCas9 RNP:LNP-mediated editing in Ai9 mice. **b.** LNP formulations used for in vivo genome editing. iGeoCas9 RNP used here was assembled from NLS-iGeoCas9(C)-2NLS and tdTom-g3(23ms). **c.** *In vivo* genome editing levels in different tissues based on GeoCas9 RNP delivery by different LNP formulations, as quantified by tdTomato signals using flow cytometry. The standard formula edits 58% of liver tissue and the cationic formula edits 35% of lung tissue. n = 6 for each group, mean ± s.e.m. **d**. Nuclei staining with DAPI (blue) and imaging of tdTomato (red) in the edited and non-edited tissues.

## Discussion

Here we describe a generalizable system for CRISPR genome editing both *in vitro* and *in vivo* based on LNP-mediated delivery of a thermostable genome editor in an RNP format. Although RNP delivery offers several potential advantages over viral-based or mRNA-based delivery strategies, its use for *in vivo* genome editing has been limited to tissue editing based on local administration/injection or liver editing through intravenous injection^20^. RNP delivery usually relies on different nanoparticle materials to encapsulate and transport RNPs; however, their applications for *in vivo* genome editing are commonly restricted by poor particle uniformity, stability and biocompatibility^48^. Use of the thermostable iGeoCas9 genome editing enzyme described here, along with newly developed LNP formulations, enables robust encapsulation and tissue-selective genome editing in mice. iGeoCas9 maintains superior stability to the commonly used SpyCas9 under a variety of conditions relevant to *in vivo* delivery and also possesses enhanced genome editing capability due to its tolerance of mutations beneficial to function while preserving molecular structure^49^. Together, LNP-mediated iGeoCas9 delivery may provide a new approach to targeted *in vivo* genetic treatments.

LNPs are a powerful non-viral delivery platform for multiple therapeutic agents including the siRNA drug patisiran and the mRNA-based COVID-19 vaccines. However, proteins are less commonly used as payloads for LNP delivery, as they tend to denature under the conditions used for LNP formulation. We hypothesized that proteins with high thermal stability and negative charge density would be efficiently delivered via LNPs because they would encapsulate readily and with preserved biochemical capability. iGeoCas9 RNP was selected as a candidate for LNP delivery because of its unique combination of thermostability, negative charge density, and genome editing functionality. Optimized LNP formulations included the synthetic pegylated lipid, ADP-2k, which, contains an acid-degradable linker that allows for rapid decomposition of the lipid in the late endosome stage of intracellular delivery, and was key to the successful development of an iGeoCas9 RNP delivery vector. The use of this LNP formulation enabled efficient iGeoCas9 RNP delivery and reduced the cytotoxicity associated with the pegylated lipid. This LNP delivery vehicle was also used for the co-delivery of RNPs and ssDNA templates to incorporate specific genomic changes by homology-directed repair. Consistent with a prior report^30^, ssDNA templates promoted RNP encapsulation into LNPs, presumably through transient binding of ssDNA to RNPs that reduced average nanoparticle size. Successful HDR corrected pathogenic mutations in disease model cell lines, highlighting the potential utility of this LNP-based delivery platform for therapeutic applications. We anticipate that this RNP:LNP strategy may be applicable to other genome-editing tools^50^ including prime editors^51^ (**Fig. S12**) and base editors^52^.

Perhaps most importantly, the LNP-assisted RNP delivery described here generated genome edits *in vivo* in both mouse liver and lung, depending on the LNP formulation used. The delivery specificity could be regulated by the electrostatic charge properties of the LNPs. In particular, LNPs prepared with 40% cationic lipid (ADC) shifted the nanoparticle targeting preference from the liver to the lung. This shift in targeting preference involves differential recruitment of plasma proteins by LNPs with different chemical properties, leading to varying cellular internalization specificity based on a receptor-mediated uptake mechanism^53^. Hints of additional tissue targeting, as evidenced by low-level genome editing in the heart, suggest that it may be possible to screen and optimize new LNP formulations to further expand delivery specificity to more challenging cell types. Together these findings demonstrate the utility of RNP:LNPs for both *ex vivo* and *in vivo* genome editing in tissues other than the liver and holds great potential for extending the therapeutic applications of CRISPR-Cas9 genomic therapy.

## Data availability

Protein, DNA, and RNA sequences in this study and next-generation sequencing data are available in the supplementary materials. Plasmid sequences, sequencing data, and raw images are available through Dryad (DOI: 10.5061/dryad.rr4xgxdfh). Other relevant data or materials are available from the corresponding author upon reasonable request.

## Acknowledgments

We thank members of the Doudna lab, the Murthy lab, and the Innovative Genomics Institute for helpful discussions. We would also like to acknowledge Ms. Netravathi Krishnappa (NGS Core Operations Manager and Sequencing Specialist, Center for Translational Genomics, Innovative Genomics Institute, UC Berkeley) for NGS and the Cystic Fibrosis Foundation Therapeutics Lab for the generous donation of 16HBEge cell lines. K.C. was supported by the Life Sciences Research Foundation. The project was funded by grants from the National Science Foundation, the Howard Hughes Medical Institute, Apple Tree Partners, and the Centers for Excellence in Genomic Science of the National Institutes of Health under award number RM1HG009490. J.A.D. is an investigator of the Howard Hughes Medical Institute. N.M. would like to acknowledge NIH grants UG3NS115599, R33 and R61DA048444-01, RO1EB029320-01A1, RO1MH125979-01, the HOPE NIH grant 1UM1AI164559, funding from the Cystic Fibrosis Foundation, the Innovative Genomics Institute, the TED foundation, and the CRISPR-Cures grant.

## Author contributions

Conceptualization: K.C., H.H., N.M., and J.A.D.; experimental studies: K.C., H.H., S.Z., and B.X.; data analysis: K.C., H.H., B.Y., M.T., and B.W.B.; supervision: J.A.D. and N.M.; manuscript writing: K.C., N.M., and J.A.D. with input from all authors.

## Materials and methods

### Plasmid construction

Plasmids used for the expression of different Cas proteins in this study were built based on a pCold vector. The inserts encoding Cas proteins contain an N-terminal CL7 tag followed by an HRV-3C protease cleavage site, and a C-terminal His_6_ tag following another HRV-3C protease cleavage sequence. The insert for the final NLS-GeoCas9(R1W1)-2NLS protein contains an N-terminal sequence consisting of different tags, His_6_-CL7-MBP (MBP: maltose-binding protein) followed by an HRV-3C protease cleavage site. The cloning reactions were carried out in a 50-μl reaction containing 1 ng of template plasmid, 1.25 μl of 10 mM dNTP, and 1.25 μl of 10 μM each primer using Phusion high-fidelity DNA polymerase (New England BioLabs). After PCR, the reactions were treated with 1 μl of DpnI (New England BioLabs) for 1 hour at 37 °C before gel purification. The plasmids were ligated based on Gibson assembly (New England BioLabs master mix) of plasmid backbone and insert sequences. The sequences of all the plasmid constructs were confirmed via full plasmid sequencing (Primordium).

### Nucleic acid preparation

All of the DNA and RNA oligos used in this study were purchased from Integrated DNA Technologies, Inc. (IDT) and HPLC or PAGE-purified. Some of the sgRNAs purchased from IDT possess chemical modifications at 3’- or 5’-ends.

### Directed evolution of GeoCas9

A chloramphenicol-resistant (CAM^+^) bacterial expression plasmid was built to have the insert gene of GeoCas9 together with its corresponding sgRNA that targets the ccdB gene in the selection plasmid with a PAM of GAAA (**g6**). Libraries of GeoCas9 mutants were generated by error-prone PCR to introduce random mutagenesis in three different regions (BH-Rec, RuvC-HNH-WED, and WED-PI). The error-prone PCR (with an error rate of 3- to 5-nucleotide mutations per kilobase) was carried out with the Taq DNA polymerase (New England BioLabs) in a reaction containing 2 µl of 10 mM primers, 1.5 µl of 10 mM MnCl_2_, 2 ng of template plasmid. The plasmid libraries were generated by ligating the mutated fragments with the remaining part of the plasmid through Gibson assembly. The plasmid libraries (∼100 ng DNA after clean-up) were electroporated into 50 µl of electronically competent cells made from *E. coli* strain BW25141(DE3) that contains the selection plasmid encoding the arabinose-inducible ccdB toxin gene. After recovery of the electroporated bacteria in 750 µl of SOB for 1.5 hours at 30 °C, the bacteria culture was concentrated; 1% of the total culture was plated onto a Petri agar dish containing only CAM (as control), and the remainder culture was plated on another Petri agar-dish containing both arabinose and CAM. Positive colonies that grew on the plates containing both arabinose and CAM were collected in a pool, retransformed (with ∼2 ng plasmid), and replated (100 µl of transformed culture on both control and selection plates). Plasmids of individual colonies from the replated plate were sequenced to obtain mutational information. Validation of the positive clones in the bacterial assay followed the same procedure.

### Protein expression

All the proteins in this study were expressed in *E. coli* BL21 (DE3) cells (Sigma-Aldrich) cultured in 2x YT medium supplemented with the antibiotics of ampicillin. The cultivation was carried out at 37 °C with a shaking speed of 160 rpm after inoculation with an overnight starter culture in LB medium containing ampicillin at a ratio of 1:40. When the optical density (OD_600_) of the culture reached 0.8–0.9, the culture was cooled down to 4 °C on ice. The expression of Cas proteins was induced by the addition of isopropyl β-D-1-thiogalactopyranoside (IPTG) to a final concentration of 0.1 mM and incubated at 15.8–16 °C with a shaking speed of 120 rpm for 14–16 hours.

To purify the Cas proteins, the cultured cells were harvested and resuspended in lysis buffer (50 mM Tris-HCl, 20 mM imidazole, 1.2 M NaCl, 10% (v/v) glycerol, 1 mM TCEP, 0.5 mM, and cOmplete protease inhibitor cocktail tablets (Millipore Sigma, 1 tablet per 50 ml) at pH 7.5), disrupted by sonication and centrifuged at 35,000 xg for 45 min. Ni-NTA resin was treated with the supernatant at 4 °C for 60 min, washed with wash buffer-1 (lysis buffer without protease inhibitor cocktail tablet), and eluted with elution buffer (50 mM Tris-HCl, 300 mM imidazole, 1.2 M NaCl, 10% (v/v) glycerol, and 1 mM TCEP at pH 7.5) to give crude His-tagged Cas proteins. The nickel elution was then subjected to Im7-6B resin in a slow gravity column repeatedly (3–4 times). The Im7-6B resin was washed with wash buffer-2 (50 mM Tris-HCl, 1.2 M NaCl, 10% (v/v) glycerol, and 1 mM TCEP at pH 7.5) before being treated with HRV-3C protease (1% weight to crude Cas protein) for 2–2.5 hours to release the Cas proteins from the CL7 and His_6_ tags. Heparin affinity column was used to further purify the desired proteins. The protein fractions were collected, concentrated, and stored in buffer (25 mM NaPi, 150 mM NaCl, and 200 mM trehalose at pH 7.50) after buffer exchange. The final yields of different Cas proteins (all with two copies of NLS at both N- and C-termini): wild-type GeoCas9, ∼10 mg per 1 L culture; GeoCas9 mutants, in a range of 2∼10 mg per 1 L culture; SpyCas9, ∼4 mg per 1 L culture; iCas12a, ∼30 mg per 1 L culture.

The purification of the final NLS-GeoCas9(R1W1)-2NLS protein is slightly different after Ni-NTA resin purification. The nickel elution was subjected to dialysis against dialysis buffer (50 mM Tris-HCl, 10 mM imidazole, 1.2 M NaCl, 10% (v/v) glycerol, and 1 mM TCEP at pH 7.5) containing HRV-3C protease (1% weight to crude Cas protein) for 12–15 hours. The tag-cleaved protein was then loaded to a heparin column and washed with 80 column volumes of buffer containing 0.1% Triton X-114 at 4 °C to minimize endotoxin impurities. The protein fractions were collected, concentrated, and subjected to further purification using a size-exclusion column in an endotoxin-free manner. The purified protein was stored in an endotoxin-free storage buffer (25 mM NaPi, 150 mM NaCl, and 200 mM trehalose at pH 7.50). The final yield of the desired GeoCas9 mutant: 5–8 mg per 1 L culture.

### Measurement of protein melting temperatures

Protein melting temperatures were measured using the thermal shift assay (GloMelt, #33021). The assay was performed on a quantitative PCR system with a temperature increase rate of 2 °C/min. The protein melting temperatures were determined as the peak values in the derivative curves of the melting curves.

### Cell lines and culture conditions

NPCs were isolated from embryonic day 13.5 Ai9-tdTomato homozygous mouse brains. Cells were cultured as neurospheres at 37 °C with 5% CO_2_ in NPC medium: DMEM/F12 (Gibco, CAT# 10565018) with GlutaMAX supplement, sodium pyruvate, 10 mM HEPES, nonessential amino acid (Gibco, CAT# 11140076), penicillin and streptomycin (Gibco, CAT# 10378016), 2-mercaptoethanol (Gibco, CAT# 21985023), B-27 without vitamin A (Gibco, CAT# 12587010), N2 supplement (Gibco, CAT# 17502048), and growth factors, bFGF (BioLegand, CAT# 579606) and EGF (Gibco, CAT# PHG0311) (both 20 ng/ml as final concentration). NPCs were passaged using MACS Neural Dissociation Kit (Papain, CAT# 130-092-628) following the manufacturer’s protocol. bFGF and EGF were refreshed every three days and cells were passaged every 5 days. Pre-coating with a coating solution containing poly-DL-ornithine hydrobromide (Sigma-Aldrich, CAT# P8638), laminin (Sigma-Aldrich, CAT# 11243217001), fibronectin bovine plasma (Sigma-Aldrich, CAT# F4759) was required for culturing cells in 96-well plates.

HEK293T and HEK293T-EGFP cells were grown in medium containing DMEM (Gibco, CAT# 10569010), high glucose, GlutaMAX supplement, sodium pyruvate, 10% FBS, and penicillin and streptomycin (Gibco, CAT# 10378016) at 37 °C with 5% CO_2_. Cells were passaged every 3 days. 16HBEge cells were grown in medium containing MEM (Gibco, CAT# 11090099), 10% FBS, and penicillin and streptomycin (Gibco, CAT# 10378016) at 37 °C with 5% CO_2_. T75 flasks pre-coated with a coating solution containing LHC-8 basal medium (Gibco, CAT# 12677-027), bovine serum albumin 7.5% (Gibco, CAT# 15260-037), bovine collagen solution, Type 1 (Advanced BioMatrix, CAT# 5005), fibronectin from human plasma (ThermoFisher, CAT# 33016-015) were used for culturing 16HBEge cells. Cells were passaged every 4‒5 days. Pre-coating was required for culturing cells in 96-well plates.

### RNP assembly

For cell culture experiments, RNPs were assembled at a 1.2:1 mole ratio of sgRNA (IDT) to Cas protein in a supplier-recommended buffer (for nucleofection) or a phosphate buffer (25 mM NaPi, 150 mM NaCl, and 200 mM trehalose at pH 7.50) immediately before use. The solution was incubated for 15–25 min at room temperature. For nucleofection, RNPs were further complexed with Alt-R Cas9 electroporation enhancer (IDT, 100-nt ssDNA) with a 1:1 mole ratio of enhancer to RNP in supplier-recommended buffers (Lonza). For LNP delivery, RNPs(±ssDNA) were further diluted with a solution of PBS/water (1:1, pH 7.3‒7.5) containing 5 mM DTT to give a final RNP concentration of 0.6‒7.5 μM.

### Genome editing with different cell lines

Nucleofection: 250k NPCs or 200k HEK293T cells were nucleofected with 100 pmol pre-assembled RNP (with 100 pmol ssDNA enhancer) with program codes of EH-100 and CM-130, respectively, according to the manufacturer’s instructions. Lonza SF (for HEK293T cells) and P3 (for tdTomato NPCs) buffers were used for the preparation of nucleofection mixtures (with a total volume of 20 µl). 10% of the nucleofected cells were transferred to 96-well plates. The culture media for NPCs was refreshed after 3 days; HEK293T cells were split with a ratio of 5:1 after 3 days. Cells were harvested for analysis after further incubation at 37°C for 2 days.

LNP delivery: 4−6.5k cells/well were seeded in 96-well plates 48 hours prior to LNP treatment (HEK293 cells: 4−5k, NPCs: 5−6k, and 16HBEge cells: 6−6.5k). The culture media was refreshed 24 hours after LNP treatment. HEK293T cells were split after 2 additional days with a ratio of 1:1 to 2:1 based on cell confluency. Cells were harvested for analysis after a total incubation time of 5 days.

### Flow cytometry

Cell fluorescence was assayed on an Attune NxT acoustic focusing cytometer (Thermo Fisher Scientific) equipped with 554 nm excitation laser and 585/16 emission filter (tdTomato), 488 nm excitation laser and 530/30 emission filter (EGFP), and 400 nm excitation laser and 440/50 emission filter (BFP). Data were analyzed using Attune Cytometric Software v5.1.1.

### Next-generation sequencing

Edited cells were harvested and treated with Quick Extraction solution (Epicentre, Madison, WI) to lyse the cells (65 °C for 20 min and then 95 °C for 20 min). Amplicons of genomic targets were PCR-amplified in the presence of corresponding primers which were designed to have no overlap with their corresponding donor ssDNA sequence in the case of HDR. The PCR products were purified with magnetic beads (Berkeley Sequencing Core Facility) before being subjected to next-generation sequencing (NGS) with MiSeq (Illumina) at 2×300 bp with a depth of at least 20,000 reads per sample. The sequencing reads were subjected to CRISPResso2 (https://github.com/pinellolab/CRISPResso2) to quantify the levels of indels and HDR.

### LNP formulations and characterization

An aqueous solution (PBS/water 1:1, with 5 mM DTT, pH 7.3‒7.5) of iGeoCas9/sgRNA RNPs (± ssDNA) was prepared and pipette-mixed rapidly with the lipid solution in ethanol at a volume ratio of 4:1 and mole ratio of 0.5‒1.1:1000, and then incubated for 1–1.5 hours at room temperature. The LNP mixture was diluted with PBS before the treatment with cell cultures. Under this procedure, RNPs were largely in excess. The size distribution of nano-formulations was measured using Zetasizer (version 7.13, Malvern Panalytical; He-Ne Laser, λ = 632 nm; detection angle = 173°).

The preparation of LNPs for animal experiments was slightly different. RNP was assembled by mixing iGeoCas9(C) and tdTom-g3(23ms) with a mole ratio of 1:1.2, incubated for 15–25 min at room temperature, and further complexed with ssDNA enhancer with a 1:1 mole ratio of RNP to enhancer. The RNP solution was further diluted with a solution of PBS/water (1:1, pH 7.3‒7.5) containing 5 mM DTT to give a final RNP concentration of 1 μM. The aqueous solution of RNP:ssDNA enhancer complexes was pipette-mixed rapidly with the lipid solution in ethanol at a volume ratio of 4:1 and weight ratio of 1:6.5 (iGeoCas9/sgRNA/ssDNA: lipids), and then incubated for 1 hour at room temperature. The LNP mixture was dialyzed against PBS using a dialysis membrane with a molecular weight cut-off of 10 kDa (ThermoFisher) at 4 °C for 2 hours and then concentrated by ultrafiltration using Amicon Ultra-15 with a molecular weight cut-off of 100 kDa (Millipore).

### *In vivo* genome editing

Two retro-orbital injections of LNPs consisting of different lipid formulations were performed with Ai9 tdTomato mice with a time window of three days. A week after the second injection, the mice were sacrificed, and all tissues were collected for further analysis. For flow analysis, isolated tissues were minced using a sterile blade and then subjected to collagenase digestion at 37 °C for 1 hour with shaking. Next, the digested solution was filtered using a 70-μm filter and quenched with PBS containing 2% FBS. A cell pellet was obtained by centrifuging for 5 min at a speed of 1500 xg at 4 °C. The supernatant was removed, and the cell pellet was resuspended in 1 ml of PBS containing 2% FBS, which could be used for flow analysis. For analysis by imaging, tissue blocks were embedded into optimal cutting temperature compounds (Sakura Finetek) and co-sectioned (8 μm) on a Cryostat instrument (Leica Biosystems) to prepare tissue sections. The mounted tissue slices were stained with DAPI before microscopy imaging. Images of tissue slices were taken using Echo Pro (v6.4.2).

